# Addition of new neurons and the emergence of a local neural circuit for precise timing

**DOI:** 10.1101/2020.03.04.977025

**Authors:** Yevhen Tupikov, Dezhe Z. Jin

## Abstract

During development, neurons arrive at local brain areas in extended period of time, but how they form local neural circuits is unknown. Here we computationally model the emergence of a network for precise timing in the premotor nucleus HVC in songbird. We show that new motor projection neurons, mostly added to HVC before and during song learning, are recruited to the end of a growing feedforward network. High spontaneous activity of new neurons makes them the prime targets for recruitment in a self-organized process via synaptic plasticity. Once recruited, the new neurons fire readily at precise times, and they become mature. Neurons that are not recruited become silent and replaced by new immature neurons. Our model incorporates realistic HVC features such as interneurons, spatial distributions of neurons, and distributed axonal delays. The model predicts that the birth order of the projection neurons correlates with their burst timing during the song.

**Significance Statement:** Functions of local neural circuits depend on their specific network structures, but how the networks are wired is unknown. We show that such structures can emerge during development through a self-organized process, during which the network is wired by neuron-by-neuron recruitment. This growth is facilitated by steady supply of immature neurons, which are highly excitable and plastic. We suggest that neuron maturation dynamics is an integral part of constructing local neural circuits.

## Introduction

During development, the birth order of neurons plays a critical role in constructing the brain’s large-scale structures. In mammalian cortex, neurons that are destined to the deep cortical layers are born earlier than those to the superficial layers [1, 2]. In rodent hippocampus, earlier born neurons and late born neurons form distinctive parallel circuits through the hippocampal pathway [3]. However, whether birth order is also important in constructing microcircuits in local brain areas is unknown [4]. The premotor nucleus HVC (proper name) of the zebra finch provides an excellent opportunity to investigate this issue.

HVC is a premotor nucleus that drives singing of the courtship song in the zebra finch [5, 6]. An adult zebra finch sings repetitions of a motif consisting of fixed sequence of syllables [7]. Excitatory HVC neurons that project to the downstream premotor area RA (robust nucleus of the arcopallium) encode the timing of acoustic features of the song [8]. Each HVC_RA_ neuron bursts once during the motif [8, 9]. As a population, HVC_RA_ neurons sequentially burst through the entire motif [10, 11].

There is strong evidence that the sequential bursting of HVC_RA_ neurons is generated within HVC [12, 9, 13, 14]. Moreover, HVC_RA_ neurons most likely form a feedforward synaptic chain network, which supports propagation of burst spikes [15, 9]. Such a microcircuit in HVC acts as an infrastructure for subsequent learning of the song, during which the connections from HVC to RA are established through reinforcement learning such that appropriate sounds are produced at appropriate time points [16, 17, 18, 19].

HVC_RA_ neurons are born and added to HVC mostly after hatching [20, 21, 22, 23]. In the zebra finch, the number of HVC_RA_ neurons almost doubles from 20 to 50 days post hatch [24], which coincides with the period of subsong and early plastic song that precede the formation of song motif. This is unlike two other major neuron types in HVC: most GABA (*γ*-Aminobutyric acid)-ergic interneurons (HVC_INT_ neurons) and neurons that project to area X (HVC_X_ neurons) are already in HVC before hatching [21] (but see [22]). Therefore, throughout song learning HVC_RA_ neurons have a wide range of birthdates.

Previous computational models [25, 26] and single unit recordings in juvenile zebra finches [27] have suggested that the feedforward synaptic chain network in HVC forms through growth by gradual recruitment of HVC_RA_ neurons to the network. However, these earlier works did not address whether the ongoing neurogenesis throughout the song learning period plays any role. Indeed, although neurogenesis in HVC has been observed for decades, its role for song learning in zebra finch has remained a mystery [28, 23].

In this paper, we propose that constant supply of newborn HVC_RA_ neurons plays a crucial role in building the synaptic chain network in HVC. We investigate this hypothesis through a computational model that builds on the previous models of network growth in HVC [25, 26]. Unlike these earlier models, our model incorporates more biologically realistic features, including explicit incorporation of HVC_INT_ neurons rather than simplifying inhibitory actions as idealized global inhibition between HVC_RA_ neurons; implementation of axonal delays between HVC_RA_ neurons, which has shown to be substantial and is important for determining the connectivity structure of the synaptic chain network [29]; and spatial structure of HVC_RA_ connectivity, which has been recently measured in zebra finch [14]. Most importantly, the maturation dynamics of HVC_RA_ neurons is modeled.

Newly born neurons have a number of properties that distinguish them from mature neurons. Immature neurons in rodents [30, 31, 32] and in songbird HVC [33] are more excitable; and in rodents, they are more amenable to synaptic plasticity [34]. In adult rodent hippocampus, these properties make adult-born dentate gyrus neurons more likely to participate in new memory formation than mature neurons [35]. We propose that newly born neurons in HVC similarly facilitate the growth of synaptic chain network. In our model, the synaptic chain network grows through spontaneous activity of neurons. Due to their high excitability, we propose that newly added HVC_RA_ neurons are preferentially recruited at the growth edge of the network. After incorporation into the network, we suggest that these neurons mature fast due to consistent activations and form a new edge of growth that leads to recruitment of a new cohort of immature neurons. This process iterates, creating a synaptic chain network that supports precise bursts of HVC_RA_ neurons. Therefore, we predict that timing of bursts correlates with the birth order of HVC_RA_ neurons during development.

We show evidence that maturity of HVC_RA_ neurons correlates with timing by reanalyzing the data from the previous experiments on juvenile zebra finch [27]. We also show that our model creates the observed spatial distribution profile for the connections between HVC_RA_ neurons [14]. With a wide delay distribution between these connections, as observed by experiments [29], our model produces a robust polychronous chain network with continuous and precise time representation, which is recently proposed to be the structure of the synaptic chain network in HVC [29]. Our model also predicts that HVC_RA_ neurons in the growing chain network receive less foreward inhibition from the HVC_RA_ neurons that drive them.

## Results

### Maturation dynamics of HVC_RA_ neurons

To investigate the possible role of immature HVC_RA_ neurons in wiring the HVC network, we created a computational model of the maturation dynamics of these neurons. We modeled HVC_RA_ neurons using two-compartmental Hodgkin-Huxley neurons with soma and dendrite (Fig. 1a), following previous models [15, 36, 9]. The somatic compartment contains sodium, delayed-rectifying potassium, and low-threshold potassium currents for generating sodium spikes. The dendritic compartment contains calcium and calcium-activated potassium currents that, in mature neurons, can generate dendritic spike that drives stereotypical tight bursts of sodium spikes in the somatic compartment.

**Figure 1:**
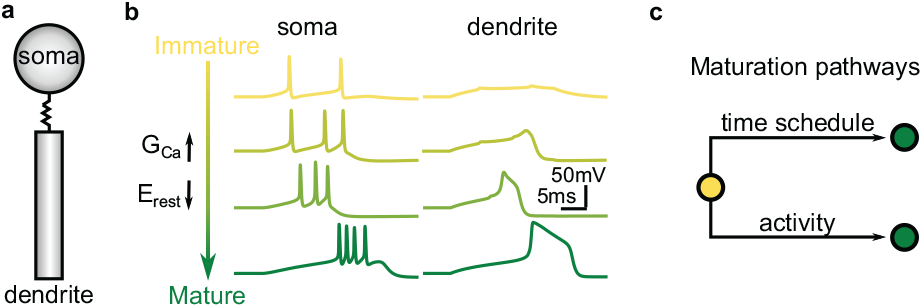
Computational model of HVC_RA_ neurons and the maturation process. (**a**) An HVC_RA_ neuron is modeled as two-compartmental Hodgkin-Huxley with soma and dendrite. (**b**) HVC_RA_ responses to the current injection to the dendritical compartment at different maturation stages. (**c**) Two pathways for neuronal maturation: scheduled maturation under spontaneous activity, and accelerated maturation driven by activity when neuron spikes reliably.

This model is modified for immature HVC_RA_ neurons. The resting membrane potential is set higher by 25 mV, since it is generally observed in rodents [31] and in HVC [33] that the resting membrane potentials of immature neurons are higher than that of mature neurons. The calcium conductance is set to zero to reflect “weak” dendritic compartment in immature neurons. Hence, immature neuron is incapable of generating tight bursts (Fig. 1b).

During maturation, the resting potential is gradually decreased and the calcium conductance is gradually increased in the dendritic compartment, eventually reaching the values for mature neurons. Dendritic calcium spike and tight burst of somatic sodium spikes gradually emerges during this process (Fig. 1b). The time course of maturation is age and activity dependent in our model (Fig. 1c). Due to elevated resting potential and noise, immature neurons spike spontaneously at ~ 0.6 Hz. A spontaneously active immature neuron matures following a time schedule, according to which both the resting membrane potential and the calcium conductance exponentially approach their mature values with time constant of 50,000 s. When a neuron is recruited into the network and spikes reliably, the maturation progressed with a faster rate, with time constant set to 500 s. In our model, spontaneous activity decreases with age, practically disappearing in adult neurons (Supplementary Fig. 8). Therefore, neurons that did not get recruited to the network gradually become silent. The silent neurons were replaced by new immature neurons in our model to mimic the continuous neurogenesis process.

### Initial HVC network

Among the three major HVC neuron types, HVC_X_ neurons have shown to have minimal impact on song production in a laser ablation study [37]. Furthermore, analysis of HVC connectivity suggests that HVC_RA_ neurons excite HVC_X_ neurons, but HVC_X_ neurons rarely connect back to HVC_RA_ neurons [38]. These results suggest that HVC_X_ are not necessary for song production. Therefore, we did not include HVC_X_ neurons in our model.

HVC of the zebra finch is roughly an ellipsoidal structure with axial dimensions 2000 *μ*m, 500 *μ*m and 500 *μ*m [14]. There are approximately 20,000 song-related HVC_RA_ neurons and 5,500 HVC_INT_ neurons [16, 39]. Due to the limitation of computational power, we could not include this many neurons in our model. Instead, we restricted ourselves to 2000 HVC_RA_ and 550 HVC_INT_ neurons. Since number of neurons is small, distributing them in 3D space becomes problematic because a large portion of them are near the boundary of the volume. To reduce this boundary effect, we placed neurons on a 2D sphere of radius 260 *μ*m. HVC_INT_ neurons were placed in a lattice-like grid on the sphere, and HVC_RA_ neurons randomly (Fig. 2a). We created connections between HVC_RA_ and HVC_INT_ neurons probabilistically according to the Gaussian distributions based on the distance between the neurons (Fig. 2b). These distributions are similar to those observed in experiments [14]. On average, an HVC_RA_ neuron connects to 65 HVC_INT_ neurons with mean distance 155 *μ*m, and an HVC_INT_ neuron connects to 115 HVC_RA_ neurons with mean distance 110 *μ*m. Initially, all HVC_RA_ neurons were immature and there were no connections between them.

**Figure 2:**
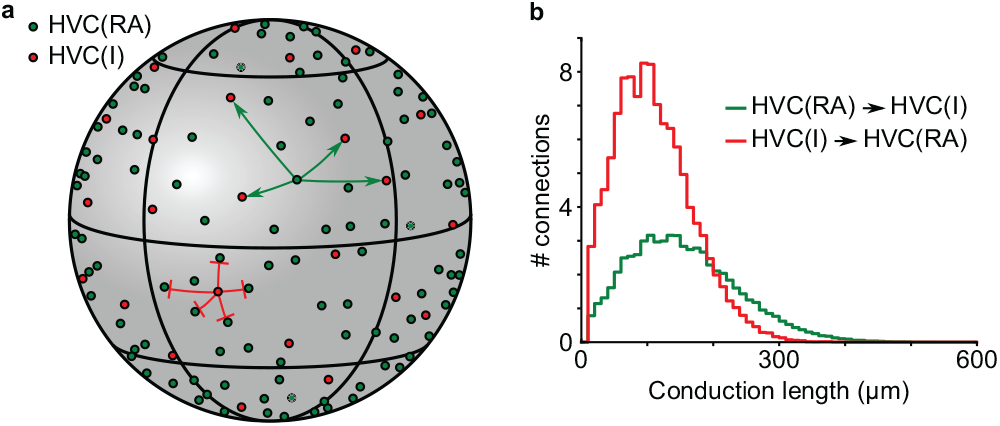
Schematic of a network arrangement and connectivity. (**a**) HVC_RA_ (dark green circles) and HVC_INT_ (red circles) neurons are distributed over the surface of a sphere. HVC_INT_ neurons form a lattice-like pattern, while HVC_RA_ neurons are distributed uniformly. Examples of connections from one HVC_RA_ neuron to HVC_INT_ neurons and from one HVC_INT_ to HVC_RA_ neurons are shown. (**b**) Distribution of axonal conduction lengths for connections between HVC_RA_ and HVC_INT_ neurons.

We also created axonal time delays between all neurons by setting the conduction velocity to 100 *μ*m/ms and using distances between neurons on the sphere. The conduction velocity was chosen such that the computed axonal delays in the model approximately match the measured axonal delays in zebra finch HVC (1 to 7.5 ms) [29].

### Growth of synaptic chain network

To grow a network of connected HVC_RA_ neurons, we used a combination of a Hebbian-like burst-timing dependent plasticity (BTDP) (Fig. 3a) and two additional plasticity rules for HVC_RA_ neurons – axon remodeling and potentiation decay, which are similar to those used in the previous models for growth of synaptic chain networks [25, 26].

**Figure 3:**
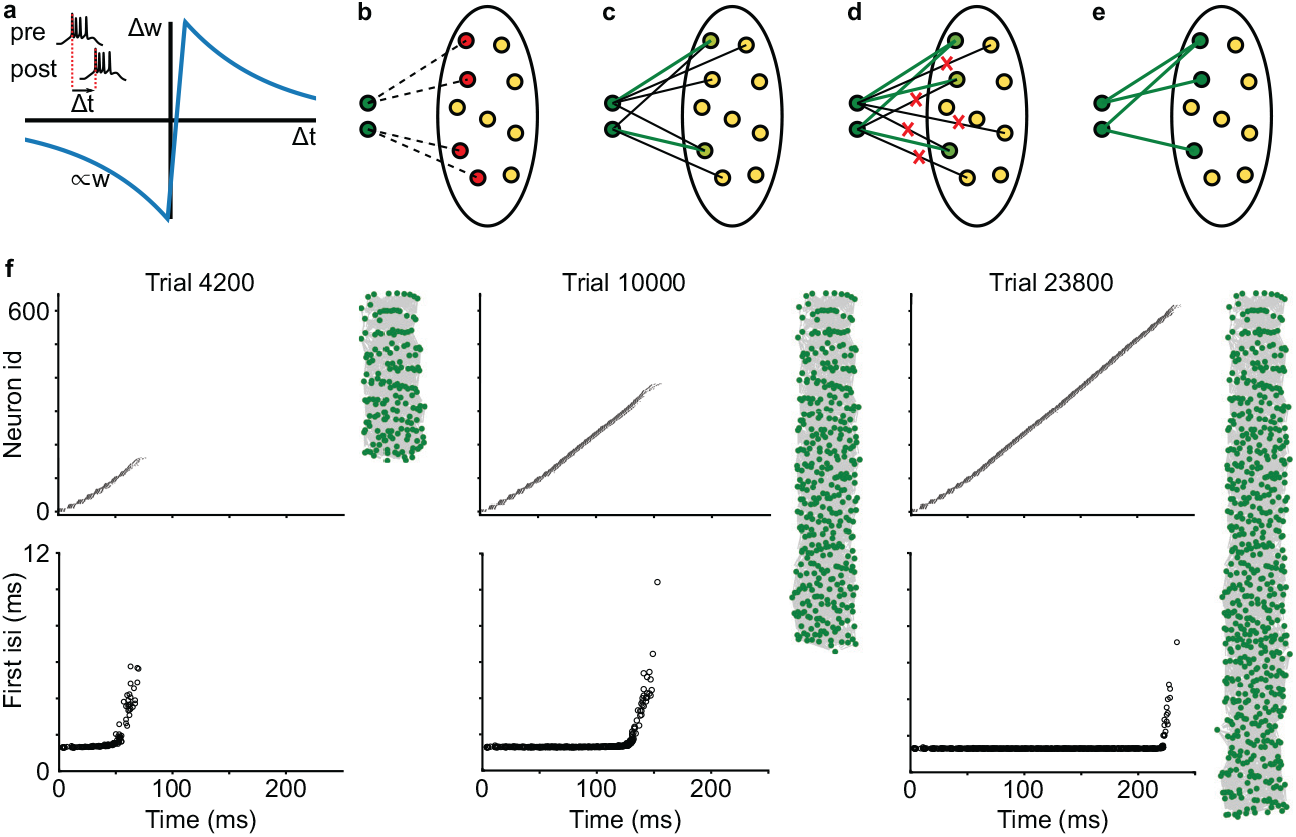
Mechanism of network growth. (**a**) Burst-timing dependent plasticity (BTDP) rule is based on the timing between burst onsets of HVC_RA_ neurons. (**b-e**) Schematic of recruitment mechanism. (**b**) Network growth begins with the starter neurons (dark green circles) activated each simulation trial and other HVC_RA_ neurons being immature (yellow circles). Silent connections (dashed lines) emerge from starter neurons to spontaneously active immature HVC_RA_ (red circles) according to the BTDP rule. (**c**) Some silent connections randomly become active (black lines), undergo further strengthening and become strong super connections (thick green lines). (**d**) When the starter neurons acquire certain number of strong super connections, other weak connections are pruned (red crosses). (**e**) The recruited neurons (dark green circles) spike reliably after the starter neurons and begin to recruit new neurons to the network. (**f**) Network growth is a gradual process in which immature HVC_RA_ neurons are added to the end of the sequence. Spike raster plots (top row) and first interspike intervals (bottom row) at different trials of the simulation are shown. Also shown are the network topology, in which green dots are neurons in the synaptic chain network and gray lines and the connections between neurons. The green dots on top are the starter neurons, and the those at the bottom are the newly recruited neurons.

BTDP was modified from spike-timing dependent plasticity rule [40]. Specifically, the time difference Δ*t* between the first spikes of the post- and pre-synaptic neurons was used, and a small positive shift was added to the time difference to ensure that no connections emerge between neurons firing synchronously. When Δ*t* > 2 ms, the synapse is potentiated (long-term potentiation, or LTP); when Δ*t* < 2 ms, the synapse is depressed (long-term depression, or LTD). The magnitude of LTP induction is maximum at Δ*t* = 5 ms, and LTD is maximal at Δ*t* = −1 ms (Fig. 3a). The magnitudes of both LTP and LTD induction decay exponentially as the absolute value of Δ*t* increases (decay constant 30 ms).

We distinguished three types of connections between HVC_RA_ neurons, depending on their strength. Silent synapses were weak, nonfunctional connections, with synaptic conductance smaller than a threshold value *W*_*a*_. They corresponded to the synapses containing only NMDA receptors [41] and did not elicit response in the postsynaptic neuron. When synaptic strength exceeded *W*_*a*_, the synapse became active and produced depolarization in the postsynaptic neuron. Strong connections with weight above *W*_*s*_ were considered as supersynaptic connections.

We randomly selected a set of 10 HVC_RA_ neurons as the training neurons, which formed a seed for the network growth. The training neurons were made fully mature with adult values for the resting potential and calcium dendritic conductance. HVC_RA_ neurons that were not in the training set, called pool neurons, started as immature neurons with high resting potential and devoid of dendritic calcium channels.

One simulation trial lasted for 500 ms in network dynamics. At each trial, the training neurons were stimulated with a synchronous kick of strong excitatory conductance. Immature pool neurons were spontaneously active during the trials due to the elevated resting potential and noise fluctuations in membrane potential. When pool neurons spiked after the training neurons, silent connections from training neurons to the pool neurons emerged according to BTDP rules (Fig. 3b). During repeating trials, silent synapses stochastically changed their strength via LTP and LTD, and can randomly became active (Fig. 3c). Emergence of too many active connections leads to uncontrolled network growth and runaway network activity. To avoid this, we introduced potentiation decay for all synapses [25, 26]. Specifically, synaptic weights of all synapses were decreased by a constant value *δ* at the end of each trial.

Depolarization of pool neurons provided by the active synapses from the training set biased these neurons to be more active during subsequent trials. Thus, a positive feedback emerged, since activity of pool neurons facilitated strengthening of synapses via LTP, eventually forming supersynaptic connections. To enforce sparse output connections, we only allowed each HVC_RA_ neuron to make a limited number of supersynaptic connections, which was set to 10 in the model. When a neuron acquired maximal number of supersynaptic outputs, the neuron underwent axon remodeling where other weak outgoing connections were pruned and did not affect their postsynaptic targets anymore [25, 26] (Fig. 3d-e). Limitations on the number of strong outputs created a competition between pool neurons for the convergent inputs from the training set. When training neurons formed the allowed number of supersynaptic connections, their postsynaptic targets were spiking reliably each iteration. The training neurons did not recruit any more targets. The recruited neurons then act as a new seed for the network growth.

In the model, network grows gradually and neurons are added to the end of the sequence (Fig. 3f). Added neurons are initially immature and have less tight burst compared to the neurons already in the sequence. With time and reliable activation, the added neurons mature and develop a tight burst. Thus, we always have immature neurons at the end of the sequence. Sequence keeps growing until all HVC_RA_ neurons are recruited into the network or its length becomes close to the length of the simulation trial.

### Axonal conduction velocity and network topology

In our model, the axonal conduction velocity controlled the axonal time delays between neurons. With the conduction velocity set to 100 *μ*m/ms, which creates the realistic axonal time delays observed in HVC [29], the emerged network showed continuous dynamics and nearly uniform temporal distribution of burst onset times (Fig. 4a). Established connections between HVC_RA_ neurons (red curve Fig. 4b) were biased towards short delay connections, but were on average longer than the preset connections to HVC_INT_ neurons. The network was temporally precise with a sub-millisecond jitter in burst onset times (Fig. 4c). Plot of the network topology based on the synaptic weights between neurons did not reveal any grouping structure (Fig. 4d). These are the characteristics of polychronous chain network proposed as the connectivity of HVC_RA_ neurons within HVC in a recent study [29].

**Figure 4:**
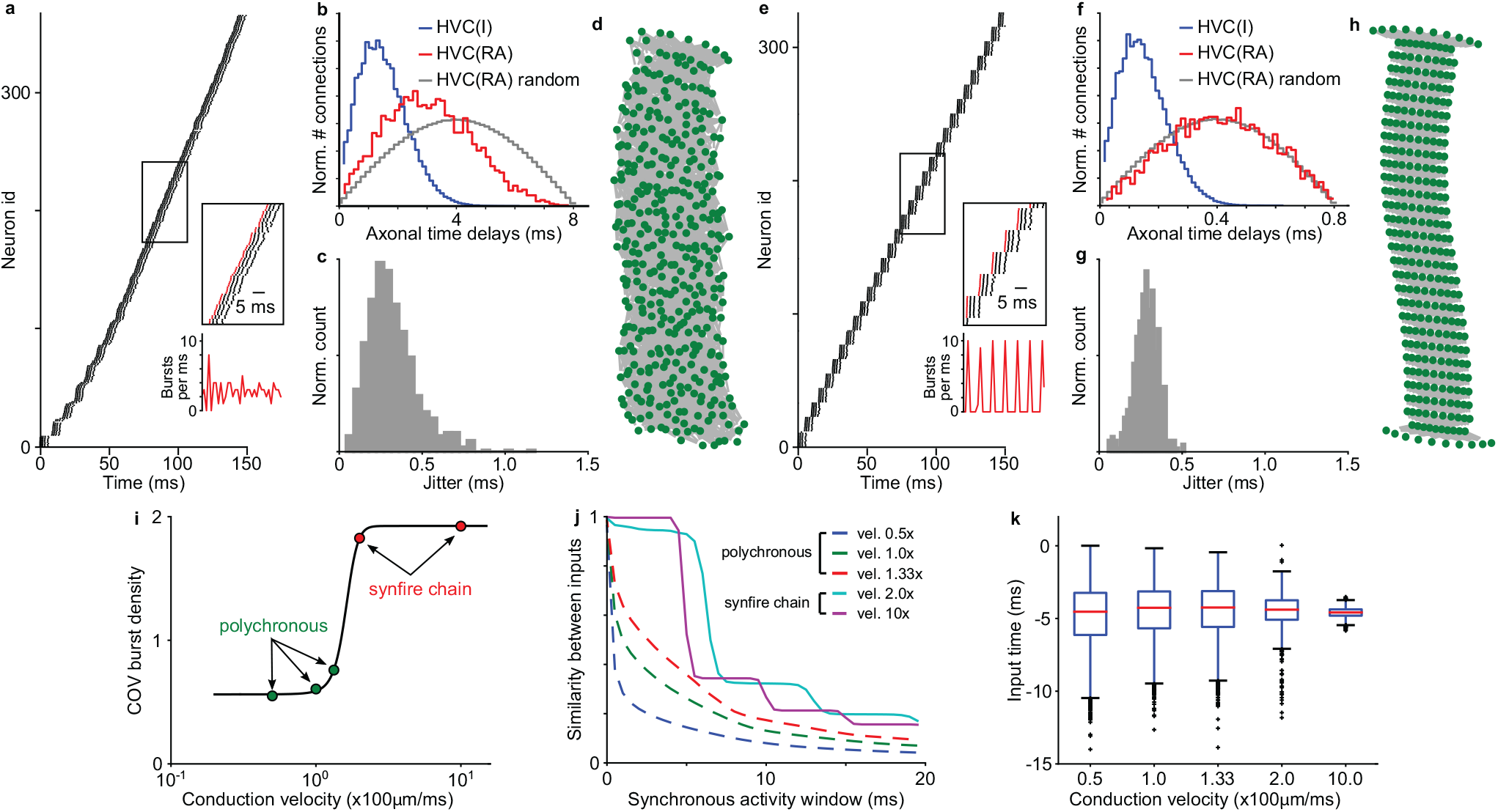
Conduction velocity shapes network topology. (**a-d**) Results for a network with conduction velocity 100 *μ*m/ms, which corresponds to the realistic axonal delays in HVC. (**a**) Raster plot of the first 150 ms of dynamics shows continuous coverage of burst onset times. (**b**) Axonal time delay distributions for efferent HVC_RA_ neuron connections to HVC_INT_ neurons (blue), formed connections to other HVC_RA_ neurons (red), and random connections to HVC_RA_ neurons (grey). Emerged connections show decrease in the number of long delay connections compared to the random connections. (**c**) Jitter in burst onset times of a grown network. (**d**) Network topology. Green dots are HVC_RA_ neurons, and the gray lines are the connections. Neurons on top are the starter neurons. Only neurons with burst onset times within first 150 ms are shown. The network has no apparent grouping of neurons. (**e-g**) Results for a network with 10x faster conduction velocity 1000 *μ*m/ms, which leads to near zero axonal delays. (**e**) Network dynamics has prominent synchronous oscillatory activity. (**f**) No bias towards shorter delay connections is observed in the grown network. (**g**) Network precision is in sub-millisecond range. (**h**) Network topology reveals groups of neurons with similar input and output connections, i.e. synfire chain layers. (**i**) Coefficient of variation of burst onset density shows transition from continuous to discrete activity pattern. (**j**) Similarity of inputs for neurons bursting within synchronous activity window has plateaus for synfire chain networks and is smooth for continuous networks. (**k**) Distributions of excitatory input times relative to burst onset time of postsynaptic neurons for different conduction velocities.

When we repeated the growth with a 10 times faster conduction velocity (1000 *μ*m/ms), the emerged network showed a strongly synchronous activity pattern (Fig. 4e). The distribution of axonal delays between HVC_RA_ neurons in the formed network was similar to the delay distribution between randomly selected pairs of HVC_RA_ neurons (Fig. 4f). The network was also temporally precise with the jitter level similar to the polychronous chain network (Fig. 4g). Network topology was highly structured, showing groups of neurons with similar input and output connections. In other words, the grown network had a synfire chain topology with prominent oscillatory activity coming from the identical chain layers of neurons.

We systematically varied conduction velocity from 0.5 to 10 times of the value measured in HVC, and observed a sharp transition in burst density oscillations at 1.5 (Fig. 4i). Networks with the velocity smaller than this value had a flat burst density, while networks with velocity exceeding this value showed prominent oscillations. We quantified the network structure using similarity of input connections for the neurons bursting synchronously in the time window of variable size (Fig. 4j). Networks with prominent oscillations in burst density (vel. 2 and 10 times) showed a stair-like decay in the similarity of inputs, which is expected for synfire chain topology with defined groups and all-to-all connections from neurons in one group to the next; whereas networks with weak activity oscillations (vel. 0.5, 1 and 1.33 times) had a smooth decreasing curve, which is expected for ploychronous chain networks with no definable groups. All grown networks, regardless synfire chains or polychronous chains, possessed a property of nearly synchronous excitatory inputs to the postsynaptic neurons (Fig. 4k).

To understand how conduction velocity influences the network topology, we examined the case of slow conduction velocity, for which the potential connections between neurons have a wide range of axonal delays. We monitored the burst onset latency of the recruited neurons relative to their presynaptic neurons (parents) (Fig. 5a). In the beginning of recruitment, connections to the recruited neurons were still weak and these neurons had a large range of burst onset latency. This permitted connections with a large range of delays to target the recruited neurons via LTP (Fig. 5b). Subsequently, however, the burst onset latency was gradually decreasing due to strengthening of the connections from the parent neurons (Fig. 5a, inset). This resulted in pruning of some of the inputs with long axonal delays via LTD (Fig. 5c). Therefore, the grown network has a prominent bias towards forming short delay connections while keeping a few long delay connections, characteristic of the delay distribution for the polychronous chain topology. In contrast, when the conduction velocity is high, all possible connections have short delays, and there is no bias towards short distance connections. In this case, synfire chain topology emerges.

**Figure 5:**
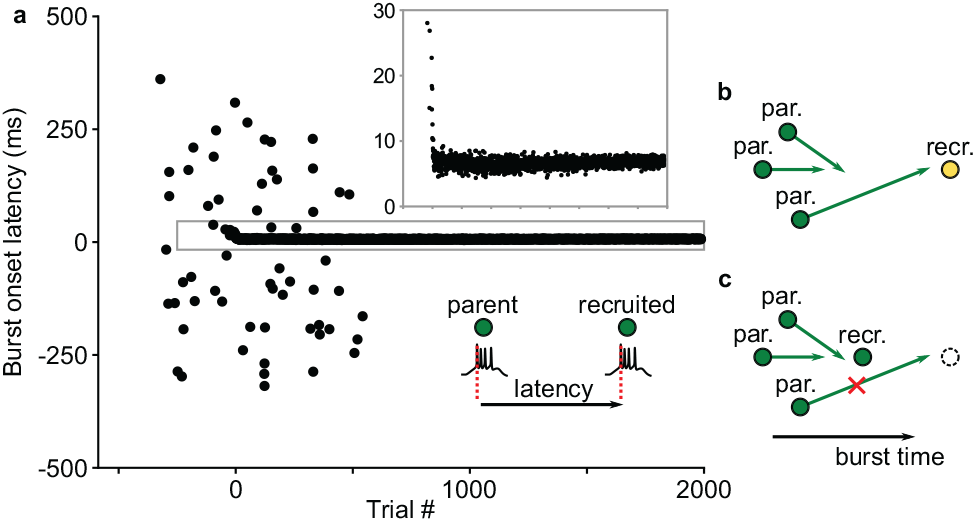
Decrease in burst onset latency of recruited neurons leads to pruning of long delay connections. (**a**) Burst onset latency between parent and recruited neurons decreases during recruitment. (**b-c**) Mechanism for pruning long delay connections. (**b**) A neuron being recruited initially spikes at a large latency, which allows long delay connections to emerge. (**c**) After recruitment, the neuron spikes at a shorter latency, which makes long delay connections to arrive late and be pruned via LTD.

### The role of inhibition in network growth

Inhibition should play an important role in network growth since it impacts the spontaneous activity of immature neurons. Due to the randomness of the connections between HVC_RA_ neurons and HVC_INT_ neurons, feedback inhibition to individual HVC_RA_ neurons is inhomogeneous in time. To see if this affects which neurons get recruited into the network, we tracked the inhibitory conductance of all HVC_RA_ neurons in the network. We considered a simulation with conduction velocity 100 *μ*m/ms (the value observed in HVC [29]) and switched off the replacement of silent non-recruited neurons to allow a direct comparison between recruited and non-recruited neurons. We observed that in the grown network, individual inhibitory connections to non-recruited neurons were stronger compared to inhibition to recruited neurons (Fig. 6a-b). Total inhibitory input, computed as a sum of all inhibitory input conductance, was also significantly larger for non-recruited neurons (*P* < 10^−42^, one-sided t-test). We then compared temporal dynamics of inhibitory conductance of recruited and non-recruited HVC_RA_ neurons during recruitment (Fig. 6d-k). When aligned to their presynaptic parent neurons (Fig. 6d-g), recruited neurons showed significantly smaller inhibitory conductance (*P* < 10^−46^, one-sided paired t-test) in LTP window, time interval which is critical for the selection of postsynaptic targets. This observation shows that neurons that receive less inhibition from the parent neurons are preferentially recruited into the growing edge of the network.

**Figure 6:**
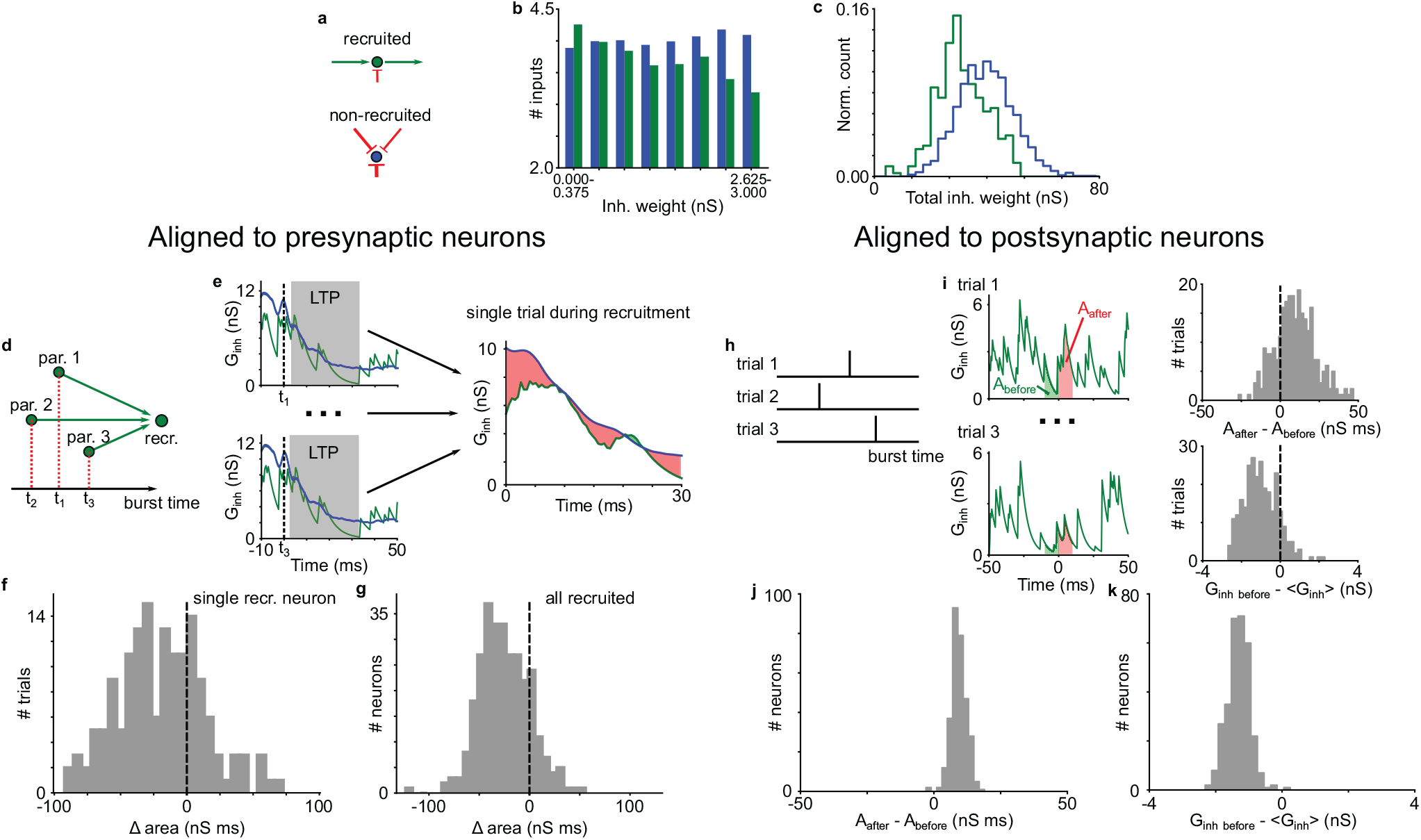
The role of inhibition in network growth. (**a-c**) Comparison of inhibitory weights onto recruited and non-recruited neurons. (**a**) Recruited neurons (green circles) receive strong excitation and weak inhibition. Non-recruited neurons (blue circles) receive strong inhibition. (**b**) Histogram of inhibitory weights shows stronger connections onto non-recruited (blue bars), compared to recruited (green bars) neurons. (**c**) Distribution of total inhibitory weights for non-recruited neurons (blue) is shifted towards stronger inhibition, compared to recruited neurons (green). (**d-g**) Comparison of inhibitory conductance aligned to presynaptic neurons during recruitment. (**d**) Inhibitory conductance is aligned to the burst onset times of presynaptic parent neurons. (**e**) Inhibitory conductance in the LTP window is averaged across all parent neurons at each trial during recruitment and compared between recruited and non-recruited neurons using the area under the conductance curve. (**e**) Difference in the area under the conductance curve for a single recruited neuron. (**e**) Difference in the area under the conductance curve for all recruited neurons. (**h-k**) Comparison of inhibitory conductance aligned to postsynaptic neurons during recruitment. (**h**) Burst times of a neuron being recruited at different simulation trials. (**i**) Inhibitory conductance is aligned to the burst onset times of recruited neurons. Difference in inhibitory conductance after and before burst is calculated using area under the conductance curve. Inhibitory conductance before burst is also compared to the mean inhibitory conductance during the trial. (**j**) Difference in inhibitory conductance after and before burst for all recruited neurons. (**k**) Difference in inhibitory conductance before burst and mean inhibitory conductance for all recruited neurons.

When aligned postsynaptically (Fig. 6h-k), recruited neurons during the recruitment show an increase in inhibitory conductance right after the burst onset time (*P* < 10^−176^, one-sided paired t-test). We attribute this observation to the self-inhibition of the neurons due to the prevalence of local connections between HVC_RA_ neurons and HVC_INT_ neurons. By bursting, HVC_RA_ neuron activated a subset of nearby interneurons, which in turn provided a feedback inhibition. The effect of such self-inhibition was not seen in the grown network due to the high network driven activity of HVC_INT_ neuron population (Supplementary Fig. 9b).

During the recruitment, the inhibitory conductance on the recruited neurons right before the burst onset time was smaller than the mean computed over the simulation trials (Fig. 6i,k, *P* < 10^−170^, one-sided paired t-test). This further supports that HVC_RA_ neurons requires less inhibition on average to be recruited. Since initial excitatory inputs to HVC_RA_ neurons are weak, the recruitment favors HVC_RA_ neurons with receiving less inhibition to ensure they can be activated by the parent neurons at the growing edge. After the network is grown and the excitatory conductance become strong, inhibitory conductance before bursts need not be small, since activations of neurons rely on strong excitatory inputs (Supplementary Fig. 9c).

### Experimental evidence linking maturity of HVC_RA_ neurons and sequence growth

The length of sequential activity of HVC_RA_ neurons grows during vocal development in zebra finches [27]. To see whether immature neurons are involved in the sequence growth, we reanalyzed the dataset of extracellular recordings in HVC of juvenile zebra finches [27, 42]. The dataset is organized into four stages of song development [27]: subsong, which is highly variable (~48 days post hatch (dph)); protosyllable song, which contains syllables with definable durations around 100 ms (~58 dph); multi-syllable song, which contains syllables with distinctive spectral characteristics (~62 dph); and motif song, which consists of a reliable sequence of syllables like adult song (~73 dph).

HVC_RA_ neurons in adult birds produce highly stereotyped bursts of 4-5 spikes lasting approximately 6 ms [8]. Experiments and computational models suggest that such a burst is driven by dendritic calcium spike [9, 15]. Since immature neurons typically do not have fully developed dendritic trees [30, 43], immature HVC_RA_ neurons may not be able to generate brief, high frequency bursts. Indeed, spike patterns of projection neurons during song development varied significantly in the number of spikes produced per burst and in the burst duration [27]. We therefore assumed that burst tightness is an indicator for HVC_RA_ neuron maturity. Specifically, we defined burst tightness as the first interspike interval in the burst (Fig. 7a). We observed that bursts in the HVC_RA_ neuron population gradually tightened as the song progressed through the protosyllable, multi-syllable and motif stages (Fig. 7b, multi-syllable versus protosyllable, *p* = 0.023, one-sided Wilcoxon rank sum test; motif versus multi-syllable, *p* < 0.0001, one-sided Wilcoxon rank sum test), supporting that burst tightness is positively linked to song development and presumably to HVC_RA_ neuron maturation.

**Figure 7:**
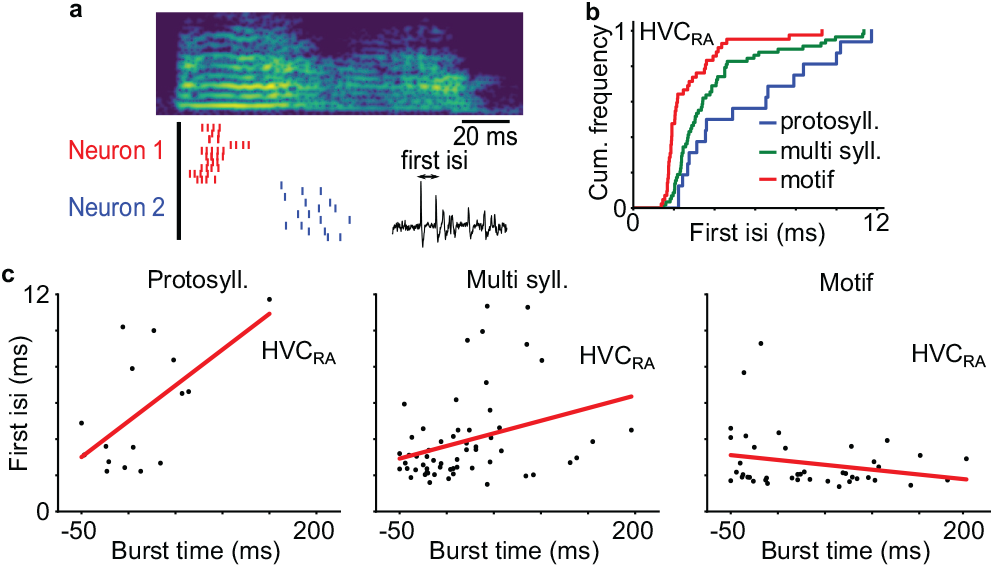
Burst tightness of HVC_RA_ neurons at different stages of songbird vocal development. (**a**) Example of spike patterns of two HVC_RA_ neurons in the protosyllable stage aligned to a syllable onset. (**b**) Cumulative distributions of first interspike intervals of HVC_RA_ neurons. (**c**) First interspike intervals of HVC_RA_ neurons at protosyllable, multi syllable and motif stages.

We next looked at the burst tightness of the HVC_RA_ neurons that are locked to syllables, *i.e.* those tend to burst at fixed latencies relative to the syllable onset times (Fig. 7c). In the protosyllable stage, the first spike interval significantly increases with the burst latency (*p* = 0.012, two-tailed t-test), suggesting that bursts are tighter for neurons bursting at the start of the syllables than those at the end. Thus, the maturity of HVC_RA_ neurons are heterogeneous in this stage, and immature neurons tend to burst towards the end of the syllables. This trend is less pronounced but still significant in the multi-syllable stage (*p* = 0.017, two-tailed t-test). It disappears in the motif stage (*p* = 0.14, two-tailed t-test).

Our analysis provides evidence that the maturity of HVC_RA_ neurons is correlated with their burst timings during song learning, and that immature neurons are preferentially added to the end of the growing sequence in HVC.

## Discussion

In adult zebra finch, HVC_RA_ neurons burst sequentially with millisecond precision during singing [8]. Electrophysiological [10] and calcium imaging [11] studies showed that the sequence is continuous, supporting the idea that such sequential bursts are generated within HVC through feedforward synaptic chain network [15, 12, 9]. Previous models suggested that such a network can be wired by recruiting neurons group by group through synaptic plasticity and spontaneous activity, resulting in growth of sequence during the wiring process [25, 26]. This prediction is in agreement with an experiment that recorded projection neurons in HVC of juvenile zebra finch [27]. Our reanalysis of this experimental data [42] suggested that HVC_RA_ neurons at the growth edge have hallmarks of immature neurons. We therefore further extended the model to include the maturation dynamics of HVC_RA_ neurons. Moreover, we included more biologically realistic features that lacked in previous models, including explicit modeling of HVC_INT_ neurons, spatial distributions of HVC neurons, and realistic axonal delays in HVC [29]. We show that immature neurons, which are more excitable hence have higher spontaneous activity rates compared to mature neurons, are preferentially recruited at the growth edge. The inclusion of the axonal delays leads to a long polychronous chain network, a structure favored by a recent analysis of HVC network and dynamics [29]. In contrast, neglecting axonal delays leads to synfire chains [44, 45], previously thought to be the topology of the HVC network [15, 25, 26]. Explicit modeling of HVC_INT_ also predicts that the wiring process favors a path of less inhibition, such that neurons that are recruited receive less forward inhibition from the recruiting neurons, highlighting the importance of inhibition in HVC [13]. Our model also reproduces the observation that HVC_RA_ neurons connect to more distal HVC_RA_ neurons, unlike their tendency to connect to nearby HVC_INT_ neurons [14].

Inclusion of immature neurons has an important effect on the growth process of synaptic chain networks. In the model, spontaneous activity plays a critical role. The distinction between immature and mature neurons allows different levels of spontaneous activity in these two populations. Immature neurons are more spontaneously active due to higher intrinsic excitability, and they are the targets of recruitments by the neurons at the growth edge. In contrast, mature neurons in the network are not spontaneously active, hence are not targets of recruitments. This allows continued growth of the network, as long as there is a supply of immature neurons in the pool. This was not the case in the previous models, in which there was a single neuron population [15, 25, 26]. There, all neurons had similar level of spontaneous activity and consequently, the chain growth usually stopped by formation of loops after neurons already into the chain were recruited. We have confirmed that loops emerge in our model as well when using a single population of mature and spontaneously active HVC_RA_ neurons (Supplementary Fig. 10).

During development, immature neurons in many neural circuits across multiple species go through a period of depolarizing inhibition before switching to hyperpolarizing inhibition, which is caused by an elevated GABA reversal potential on immature neurons [46]. Our computational experiments with developmental switch in GABA resulted in the emergence of numerous connections between nearby HVC_RA_ neurons (data not shown). This was because dense local connectivity between HVC_RA_ and HVC_INT_ neurons promotes recruitment of nearby immature neurons through depolarizing local inhibition. Experimentally, local connections between HVC_RA_ neurons are sparse [14]. We therefore assumed that the emergence of connectivity between HVC_RA_ neurons happens at the time when GABA exerts an adult hyperpolarizing response on immature neurons. This assumption needs to be tested in future studies with intracellular recordings of HVC_RA_ neurons during development.

In our model, maturation of immature neurons is activity driven. Spontaneously activity alone is enough for the neuron to mature, but more reliable activation after recruitment into the network accelerates the maturation. This acceleration protects the grown network from spontaneous activation and hence from formation of loops. This maturation dynamics is inspired by the observation in rodent hippocampus that adult-born neurons mature faster with enhanced activity and mature more slowly with reduced activity [47]. The exact value of the activity-driven maturation time scale is not important, as long as it is much smaller than the spontaneous one. Neurons that become mature but not recruited into the network become silent eventually and are replaced by a fresh immature neuron. This turnover ensures that there is a fresh supply of immature neurons for the chain growth. The rate of replacement also controls the number of available targets for the growth, which is important for forming convergent inputs to the targets during the recruitment process. If the number of targets is too large, recruiting neurons can connect to divergent targets, and the resulting network is not capable of producing precise timing. A consequence of the turnover is that the bursting timing of neurons in the chain network is positively correlated with the order of their introduction. In other words, timing correlates with birth order. This prediction of our model can be tested by labeling cohorts of newborn neurons using viral strategy in juvenile [22] and recording their burst timings in adulthood using calcium imaging [11].

Addition and turnover of HVC_RA_ neurons post hatch has been observed for over 30 years [20, 48], but the significance of this process for birdsong learning and production remains unclear [28, 23]. In juvenile zebra finch, deprivation of auditory inputs by deafening before song learning [49] and inability to learn tutor song due to peripheral nerve injury [50] did not impact recruitment of HVC_RA_ neurons. These observations are consistent with our view that addition of HVC_RA_ neurons mainly contributes to the self-organized wiring process of the synaptic chain network in HVC, which should not depend on auditory inputs or learning specific tutor song.

Synfire chain is a popular feedforward model generating precise and stable sequential activity of neurons [44, 45, 51]. Several computational models have explored the formation of synfire chains. Successful models that can grow long sequences use a combination of STDP rules and additional synaptic plasticity mechanism to constrain the connectivity. With STDP rule and heterosynaptic plasticity rules that limit the total incoming and outgoing synaptic weights for each neuron, Fiete et al [52] showed formation of synfire chain loops with length distributed according to a power law. Short loops were more numerous than long loops. However, to form groups of neurons that fire at the same time as observed in HVC, the model needed to introduce additional correlated inputs that defined coherent groups before chain formation. Jun and Jin showed that synfire chain forms with Hebbian STDP and additional synaptic plasticity rules that constrain the number of strong output connections. The model was able to show the gradual growth of synfire chains through group-by-group recruitment of HVC_RA_ neurons. The process ends with the formation of a loop, with length following a Gaussian distribution [26].

Our study builds upon the gradual recruitment model [25, 26] and uses similar synaptic plasticity rules. However, our model introduces several realistic features that none of the previous models had, including explicit modeling of HVC_INT_ neurons; spatial distributions of neurons and realistic axonal time delays recently measured in HVC [29]; and, most importantly, newly born HVC_RA_ neurons and their maturation dynamics. These lead to novel insights, as discussed earlier. Additionally, no loops form in our model, unlike all previous models. Under realistic axonal time delays, we show that a continuous polychronous network rather than synfire chain emerges after the training. The network still possesses a sub-millisecond level of precision and its burst times cover the sequence almost uniformly with no silent gaps. We also show that by using connections with fast conduction velocity, we can recover the synfire chain topology. Grown synfire chain has similar sub-millisecond level of precision, but its burst density shows prominent oscillations. We demonstrate that by changing axonal conduction velocity between HVC_RA_ neurons, we can grow either synfire chain or polychronous chain network. In the polychronous chains, neurons are driven by almost synchronous inputs despite of distributed presynaptic spike times due to the delays. This is similar to a previous study in which approximately 70 ms long polychronous sequences with an average size around 20 neurons emerged and disappeared in a recurrent network with STDP rules for synaptic plasticity [53]. However, in our case incorporation of additional synaptic plasticity rules produce stable sequences that span hundreds of milliseconds and contain hundreds of neurons. Thus, we show that long polychronous neuronal sequence can emerge from a combination of STDP and additional synaptic plasticity rules.

Our growth algorithm is robust with respect to the changes in the model parameter values. The use of different strength of inhibitory connections (varied between *G*_*ie*_ = 0.015 *mS*/*cm*^2^ and *G*_*ie*_ = 0.060 *mS*/*cm*^2^), different number of efferent supersynaptic connections (*N*_*s*_ = 10 and *N*_*s*_ = 20), and different maximal strength of excitatory connections between HVC_RA_ neurons (between *G*_*max*_ = 1.5 *nS* and *G*_*max*_ = 4 *nS*) lead to the emergence of precisely timed neural sequences (data not shown). Thus our modeling results do not rely on fine-tuning of the model parameters.

Our re-analysis of the data that recorded HVC neurons in juveniles [27, 42] showed that burst tightness of projection neurons decreases with the burst timing during the sequence growth in the protosyllable state. This difference disappears in later stages of song learning. We interpreted the less tightness of bursts as a reflection of immature intrinsic bursting mechanism. An alternative possibility is that the burst tightness is a network phenomenon. It is possible that neurons that burst earlier in the sequence are better connected and get stronger inputs, leading to tight bursts, whereas those that burst later are still in process of getting incorporated and hence are loosely connected. Another possibility is that feedback inhibition controls the burst tightness [54]. There is some evidence in the data that supports the intrinsic mechanism. We found one HVC_RA_ neuron in the subsong stage that was not locked to vocalization but still showed tight bursts usually observed in the motif stage (Supplementary Fig. 11). Since the network is unlikely formed in this stage, this observation favors intrinsic mechanism for burst tightness. Due to limited number of HVC_RA_ neurons recorded in subsong stage and subsequent protosyllable stage, we could not gather more evidence. Future experiments with more data on HVC_RA_ neurons in early song learning stages, perhaps also including intracellular recordings *in vivo* and in slices, should be able to address whether burst tightness is intrinsically controlled.

We use synaptic plasticity rules based on the timing of burst onsets (BTDP). This simple rule sidesteps the complex interaction of multiple spikes within the bursting pre- and post-synaptic neurons [55], and is guided by the observation that in cortical neurons, the timings of the first spikes in bursts are most important for determining the timing-dependent LTP and LTD [56]. In addition, we apply a small 2 ms shift of BTDP curve to the region of positive times, so that there is an LTD for synchronously bursting neurons. This prevents the emergence of connections between neurons that fire synchronously. Such a shift was used to stabilize weight distributions in random networks of spiking neurons in another modeling study [57]. Whether these rules apply to synaptic plasticity for HVC_RA_ neurons remains to be seen. To date, there is no systematic study of synaptic plasticity in HVC, and further experiments are needed.

In addition to sequence growth, extracellular recordings in juvenile zebra finches also revealed sequence splitting during the syllable development [27]. At the protosyllable stage, majority of the projection neurons fired in a single protosequence. When several syllable types emerged from a common protosyllable, the corresponding protosequence split. While there were still neurons firing at all syllables with the same latencies relative to syllable onsets (“shared neurons”), more neurons fired specifically to a single syllable type. Gradually, the shared neurons disappeared. The authors proposed a model, according to which a protosequence grown from a common seed of synchronously activated neurons is split by dividing the seed into several groups activated at different times, and also by increasing local inhibition. In our study, the splitting does not happen during the network growth and we did not explore mechanisms for it to happen. Activation of seed neurons at different times and increase in inhibition may also induce protosequence splitting in our model.

In conclusion, we have shown that protracted addition of new neurons in HVC in juvenile helps to wire synaptic chain network through a self-organized process. Our model illustrates the possibility that birth order of neurons is important for constructing functional microcircuits in local brain areas.

## Methods

### Juvenile zebra finch data analysis

We reanalyzed a previously reported data set of extracellular recordings in HVC of juvenile zebra finches [27, 42]. The data set contained recordings of projection neurons from 32 birds during the song development (44-112 dph). HVC_RA_ neurons exhibited sparse bursting activity. Following the procedure in Okubo et al [27], a burst was defined as a continuous group of spikes separated by intervals of 30 ms or less. To determine the burst tightness of a projection neuron, we estimated the median of the first interspike intervals of all the bursts produced by the neuron at a given song learning stage (subsong, protosyllable, multi-syllable, and motif). To find the bursting time of the neurons locked to syllables, we followed the approach in Okubo et al [27].

### Network model

We distributed 2000 HVC_RA_ and 550 HVC_INT_ neurons over the 2-D sphere of radius 260 *μ*m with no overlap. A neuron occupies a volume of a sphere with diameter 10*μm*. HVC_INT_ neurons were first placed evenly on the sphere using the Fibonacci lattice [58]. The distance between nearest neighbors on sphere is approximately Δ*r*_*in*_ = 40 *μ*m, which matches the average distance between HVC_INT_ in real HVC (as estimated from the HVC volume and the number of interneurons). Then, they were randomly shifted along the sphere surface by a small amount: Δ*θ* = 0.0006Δ*r*_*in*_ and Δ*ϕ* = 0.0006Δ*r*_*in*_/*sin*(*θ*), where *θ* is the latitude of a neuron’s position on the sphere, *ϕ* is its longitude. HVC_RA_ neurons were placed randomly over the surface sphere, with the constraint that they do not overlap with other HVC_RA_ or HVC_INT_ neurons.

Connections between HVC_INT_ and HVC_RA_ neurons were placed probabilistically based on the distance between neurons along the sphere: 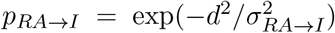 and 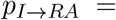 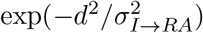, where *p*_*RA*→*I*_ is a probability for a given HVC_RA_ neuron to contact a given HVC_INT_ neuron, *p*_*I*→*RA*_ is a probability for a given HVC_INT_ neuron to contact a given HVC_RA_ neuron, *d* is a distance between given HVC_RA_ and HVC_INT_ neurons on the sphere, *σ*_*RA*→*I*_ = 130 *μm*, and *σ*_*I*→*RA*_ = 90 *μm*. Only a single connection between a pair of neurons was allowed. Parameter *σ*_*RA*→*I*_ was chosen to match the upper bound on the number of postsynaptic HVC_INT_ partners for an HVC_RA_ neuron [14, 59]. On average an HVC_RA_ neuron contacted 11.6% of HVC_INT_ neurons. HVC_INT_ neurons had a smaller spatial connectivity scale to influence nearby HVC_RA_ neurons. A single HVC_INT_ neuron contacted 5.8% of HVC_RA_ neurons. Conductance of the connections were sampled from uniform distributions on the intervals (0, *G*_*ei*_) for HVC_RA_ to HVC_INT_ connections and (0, *G*_*ie*_) for HVC_INT_ to HVC_RA_ connections, with *G*_*ei*_ = 0.4 *mS*/*cm*^2^ and *G*_*ie*_ = 0.03 *mS*/*cm*^2^. Axonal time delays for the connections were calculated by multiplying the distance between neurons by axonal conduction velocity. Normal conduction velocity was set to 100 *μ*m/ms, as observed in HVC [29]. Connections between HVC_RA_ neurons did not exist at the start of simulations.

A randomly selected set of 10 HVC_RA_ neurons were chosen as the starting seed for the network growth. The training neurons had the mature properties, while other HVC_RA_ neurons started as immature.

### Growth simulation

Network dynamics was run in trials of 500 ms duration with a time step 0.02 ms. In the beginning of each trial, the dynamical variables of neurons were reset to their resting values. At a random time between 100 ms and 400 ms in trial, the training neurons were excited by a synchronous excitatory conductance kick of strength 300 nS, which made them burst. Simulations were run until the number of supersynaptic connections in the network remained constant for 10000 trials.

### Neuron model

For HVC_INT_ neuron we used a single compartment Hodgkin-Huxley model identical to the one described in [9]. For HVC_RA_ neuron we used a two-compartmental Hodgkin-Huxley model with soma and dendrite similar to the one in [9].

Parameters of sodium, potassium and leak currents of the soma of a mature HVC_RA_ are identical to those in [9]. Somatic compartment is additionally equipped with low-threshold potassium current *I*_*KLT*_ = *G*_*s,KLT*_*l*(*V*_*s*_ − *E*_*K*_) with conductance *G*_*s,KLT*_ = 3.5 *mS*/*cm*^2^, potassium reversal potential *E*_*K*_ = −90 *mV* and gating variable *l*. Gating variable obeys the following dynamics: *τ*_*l*_*dl*/*dt* = *l*_∞_(*V*) − *l*, where *τ*_*l*_ = 10 ms, *l*_∞_(*V*) = 1/(1 + exp −(*V* + 40)/5). Parameters of the dendritic compartment of a mature HVC_RA_ are identical to [9], except for *τ*_*c*_ = 15 ms.

Immature HVC_RA_ neuron has elevated leak reversal potential *E*_*L*_ = −55 *mV* in both somatic and dendritic compartments. In addition, the calcium conductance in the dendritic compartment of immature HVC_RA_ were set to zero.

### Synapse model

Synaptic conductances on neurons were modeled according to “kick-and-decay” dynamics [9]. Synaptic conductance of a neuron increases following a delivery of a spike to the synapse with conductance G: *g*_*syn*_ → *g*_*syn*_ + *G*. In between spike arrivals, synaptic conductance decays exponentially: *τ*_*syn*_ *dg*_*syn*_/*dt* = −*g*_*syn*_. We used the same values for synaptic decay time constants as in [9].

### Noise model and simulation

Noise in HVC_INT_ neurons was created using stochastic Poisson spike trains arriving at excitatory and inhibitory synapses, mimicking random synaptic activity, such that HVC_INT_ neurons spiked spontaneously with rate ~ 10 Hz. Parameters of the Poisson spike trains were identical to [9]. Dynamics of HVC_INT_ neuron was solved using Dormand-Prince order 8 method [60].

Noise in HVC_RA_ neurons was implemented by injecting white noise current of amplitude 0.1 nA to soma and 0.2 nA to dendrite [29]. To account for white noise stimulus, HVC_RA_ model was treated as a system of stochastic differential equations and was solved with weak order 3 AN3D1 method [61].

### Maturation model

Maturation of HVC_RA_ neurons was modeled as a gradual increase of dendritic calcium conductance, and a gradual decrease in the somatic and dendritic leak reversal potential:

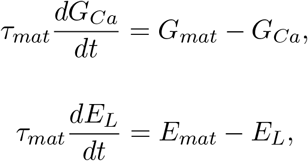

where *τ*_*mat*_ is the maturation time constant; *G*_*mat*_ = 55 *mS*/*cm*^2^ is the mature value of calcium conductance; and *E*_*mat*_ = −80 *mV* is the mature value of leak reversal potential. Values of *G*_*Ca*_ and *E*_*L*_ were updated at the end of each trial. Maturation rate of an HVC_RA_ neuron *τ*_*mat*_ depended on its activity history. If a neuron spiked in less than half of the trials in the past 1000 trials, it was treated as spontaneously spiking. Once a neuron spiked in more than half of the trials in the past 1000 trials, it was treated as reliably spiking. For a spontaneously spiking neuron, maturation time constant was set to *τ*_*mat*_ = 50,000 s. For a reliably spiking neuron, maturation time constant was set to a smaller value of *τ*_*mat*_ = 500 s.

### Neuronal turnover

Neuron was assigned as silent if it spiked in less than 80 trials in the past 4000 trials. Silent neurons were replaced at the end of each trial with immature neurons. New immature neurons were placed randomly on the surface of the sphere representing HVC, avoiding overlaps with all HVC_RA_ and HVC_INT_ neurons.

### BTDP synaptic plasticity rule

To update weights between HVC_RA_ neurons, we used a BTDP rule based on burst onset timing between presynaptic and postsynaptic neurons (Fig. 3a). We defined a “burst” as a continuous group of spikes with duration 30 ms or less. Burst onset time was defined as the first spike in a burst. Each time a neuron produced a new burst, all afferent synapses onto the neuron and all efferent synapses are updated. For a pair of a presynaptic neuron *i* with burst onset time *t*_*i*_ and a postsynaptic neuron *j* with burst onset time *t*_*j*_, an additive LTP would occur for the synapse with weight *G*_*ij*_ if Δ*t* = *t*_*j*_ − *t*_*i*_ > *T*_0_:

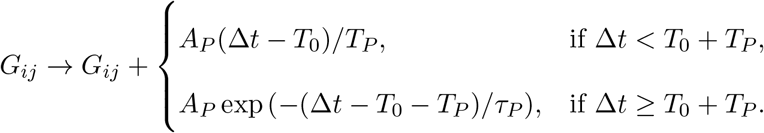

If Δ*t* ≤ *T*_0_, the synapse undergoes depression through multiplicative LTD:

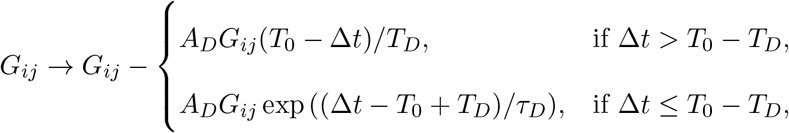

The following parameters were used in simulations unless specified: *A*_*P*_ = 0.25 nS, *A*_*D*_ = 0.02, *T*_0_ = 2 ms, *T*_*P*_ = 3 ms, *T*_*D*_ = 3 ms, *τ*_*P*_ = 30 ms, *τ*_*D*_ = 30 ms. All weights were clipped below *G*_*min*_ = 0 nS and above *G*_*max*_ = 4 nS.

### Synapse states

Synapses were in 1 of 3 possible states depending on their synaptic weight. Synapses with weights 0 < *W* < *W*_*a*_ were silent and did not elicit response in postsynaptic neurons. Synapses with weights *W*_*a*_ < *W* < *W*_*s*_ were active and produced depolarization in postsynaptic neurons. Synapses with weights *W* > *W*_*s*_ were supersynapses that produced a strong response in postsynaptic neuron. Regardless of their state, all synapses participated in BTDP update rules. The following parameters were used in simulations unless specified: *W*_*a*_ = 0.2 nS, *W*_*s*_ = 1.0 nS.

### Potentiation decay

All synapses experience a depression at the end of each trial: *G* → *G* − *δ*, where *δ* = 0.01 nS. This depression is needed to prevent the emergence of too many active synapses that may lead to uncontrolled network growth [26].

### Axon remodeling

The axon remodeling rule was identical to the one in [25]. When the number of efferent supersynaptic connections of a neuron reaches *N*_*s*_ = 10, the neuron is saturated and all other active efferent connections of the neuron are withdrawn. Withdrawn connections do not elicit effect on postsynaptic neurons and do not participate in BTDP updates. However, they still undergo potentiation decay. Withdrawn connections will be re-connected if the neuron loses one or more of its supersynapses.

### Neural activity analysis

Burst density was calculated as a histogram of burst onset times with bin size 1 ms. The presence of oscillations in burst density was estimated using the coefficient of variation (CV), which is a standard deviation divided by the mean. Jitter in a neuron’s timing was calculated as a standard deviation of the burst onset times based on the 200 test runs of the dynamics of the grown network.

### Network structure

Plots of network topology were based on the supersynaptic weights between neurons and were created using Kamada-Kawai algorithm in Pajek software program for network analysis [62].

Network structure was also analyzed using the similarity of inputs to neurons that spike synchronously within a time window *T*_*w*_. For neuron *i* that bursts at *t*_*i*_, the synchronously spiking neurons have their burst onset times within a time interval (*t*_*i*_ − *T*_*w*_/2, *t*_*i*_ + *T*_*w*_/2). The similarity of inputs to neuron *i* and a synchronously spiking neuron is computed as the fraction of the presynaptic neurons common to the two neurons among all presynaptic neurons to the two neurons (the Jaccard index). The mean Jaccard index of all synchronously spiking neurons at *t*_*i*_ represents the similarity of inputs at this time. The mean Jaccard index for all burst times is defined as the similarity of inputs for a given time window *T*_*w*_.

### Analysis of inhibition

With neuronal turnover disabled and the conduction velocity set to 100 *μ*m/ms, inhibitory conductance of all HVC_RA_ neurons was tracked for 30000 trials. By the end of these trials, the number of supersynaptic and active connections have reached stable values and the network growth stopped. A neuron was designated as recruited if it spiked consistently during the testing trials of the grown network in more than 95 out of 100 trials. The time of its recruitment was estimated using its spike history during the growth. At each trial, the number of the neuron’s spikes averaged over a window of the past 25 trials was computed, and when the average first reached 1, which signaled the start of reliable spiking, the trial was defined as the trial at which the neuron was recruited.

For a recruited neuron *i*, an LTP window is defined relative to the burst time of its presynaptic neuron *j*, during which the synaptic strength from neuron *j* to neuron *i* can be strengthened according to the BTDP synaptic plasticity rule. Specifically, the window is the time interval (*t*_*j*_ + *d*_*ji*_ + *T*_0_, *t*_*j*_ + *d*_*ji*_ + *T*_0_ + *τ*_*P*_), where *d*_*ji*_ is the axonal delay; *T*_0_ = 2 ms is the time shift in BTDP synaptic plasticity rule; and *τ*_*P*_ = 30 ms is the time scale of the LTP part of BTDP. At each trial before the recruitment, a set of inhibitory conductance traces on neuron *i* is extracted in the LTP windows relative to all its presynaptic neurons. The average of this set represents an inhibitory conductance of the recruited neuron at trial *T* aligned to its presynaptic neurons.

For comparison, an average inhibitory conductance of non-recruited neurons is extracted in the same time intervals, and is defined as the inhibitory conductance of non-recruited neurons. Difference in the area under conductance curves is computed numerically using a trapezoid method. The median difference in the area computed for all trials before the recruitment represents the difference in the inhibitory conductance between the recruited neuron and the non-recruited neurons.

For analysis of inhibition on a recruited neuron *i* relative to its burst onset times before the recruitment, only trials in which neuron *i* produced bursts are considered. For each such trial, the area under the inhibitory conductance curve is calculated for 10 ms before and 10 ms after the burst onset time. The median difference in area for all trials represents the difference in the inhibitory conductance before and after bursting of neuron *i*. The difference of the inhibitory conductance before burst relative to the average is defined as median of the differences between the mean inhibitory conductance 10 ms before the burst and the mean during the trial for all trials before the recruitment.

To investigate the inhibition after recruitment, similar procedure is applied to 100 test trials of the grown network.

## Acknowledgements

Research supported by NSF award EF-1822476, the Huck Institutes of the Life Sciences at Pennsylvania State University, and generous donation from Indus Capital in memory of Shunyu Zheng. We thank Ben Scott and Michael Long for useful comments on the manuscript.

## Supplementary Figures

**Figure 8:**
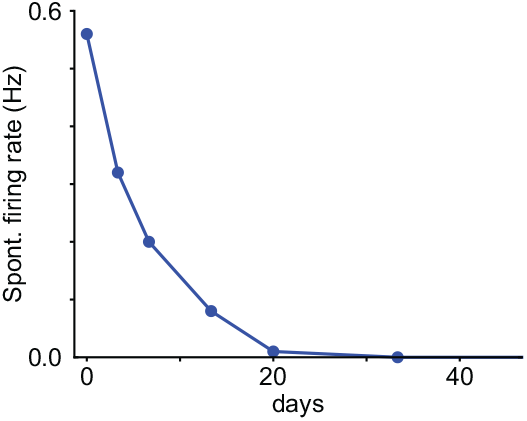
In the model, spontaneous firing rate of HVC_RA_ neuron decreases with neuronal age due to reduced excitability.

**Figure 9:**
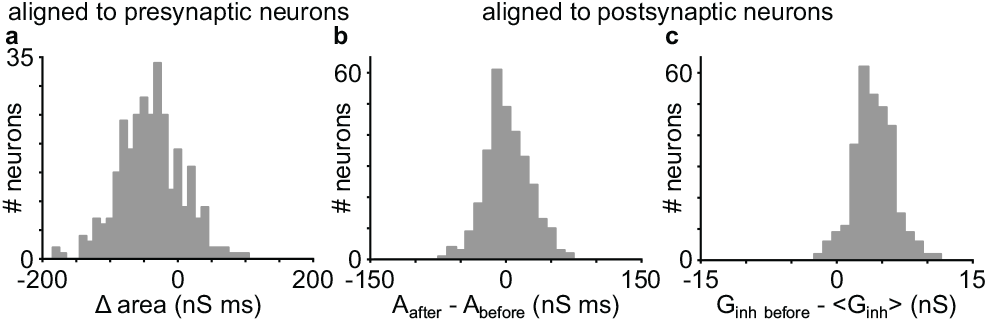
Comparison of inhibitory conductance for a grown network based on 100 test trials. (**a**) Difference in the area under the conductance curve in the LTP window for all recruited neurons aligned to presynaptic parents. (**b-c**) Analysis of inhibitory conductance of recruited neurons aligned postsynaptically. (**b**) Difference in inhibitory conductance after and before burst for all recruited neurons. (**c**) Difference in inhibitory conductance before burst and mean inhibitory conductance for all recruited neurons.

**Figure 10:**
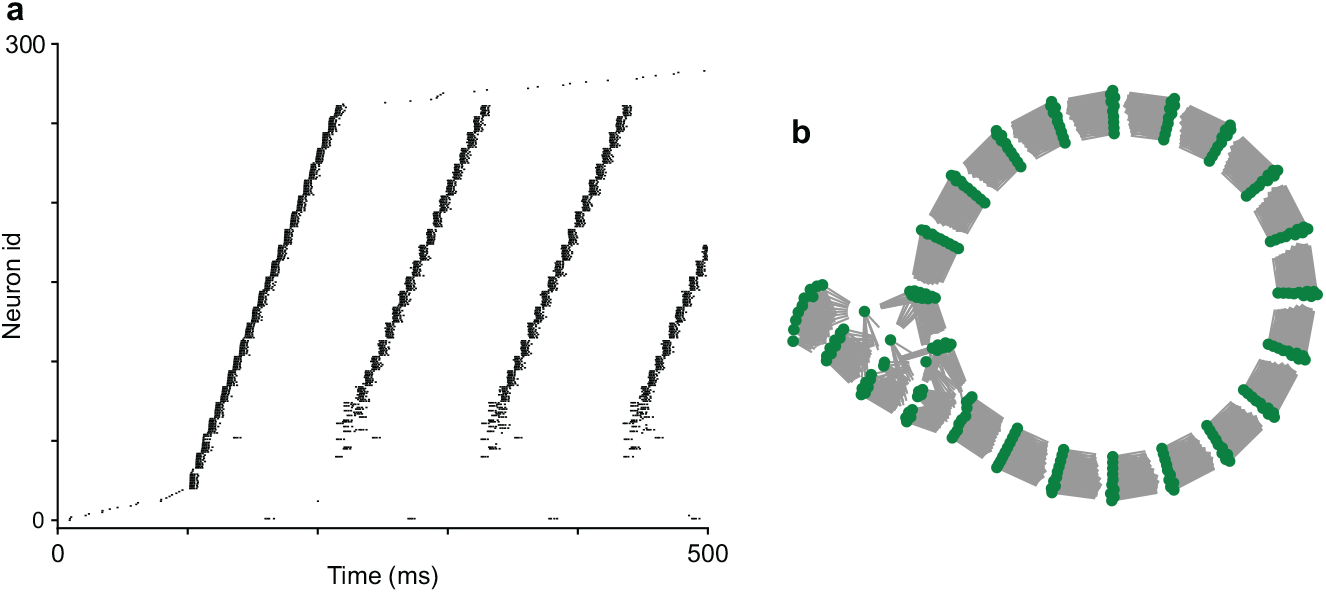
Loop formation in the network with noisy mature HVC_RA_ neurons. When we use a single population of mature spontaneously active HVC_RA_ neurons receiving a large white noise stimulus of amplitude 0.25 nA to soma and 0.5 nA to dendrite, loop sequences form. Here we use a fast conduction velocity 1000 *μ*m/ms, which leads to the emergence of a synfire chain. (**a**) Raster plot of network dynamics. (**b**) Network topology based on synaptic weights between neurons.

**Figure 11:**
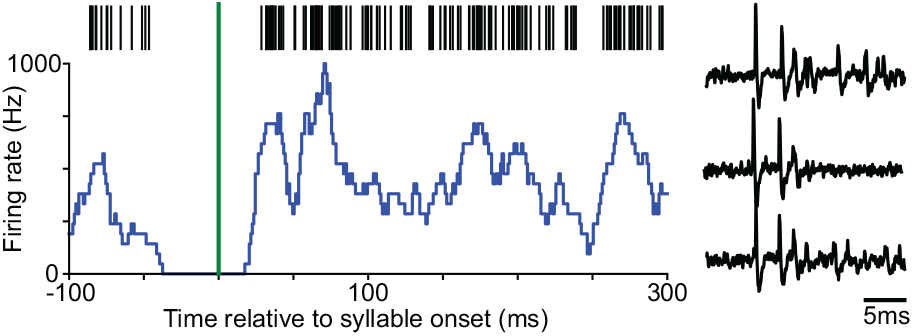
Example HVC_RA_ neuron recorded in the subsong stage showing tight burst without being locked to the song. (Left) Firing rate of the neuron aligned to syllable onset times does not show significant peak, meaning that the neuron is not locked to the syllables. (Right) Example membrane potential traces of the same neuron demonstrate tight bursting pattern.

